# Easy quantification of template-directed CRISPR/Cas9 editing

**DOI:** 10.1101/218156

**Authors:** Eva K. Brinkman, Arne Nedergaard Kousholt, Tim Harmsen, Christ Leemans, Tao Chen, Jos Jonkers, Bas van Steensel

## Abstract

Template-directed CRISPR/Cas9 editing is a powerful tool for introducing subtle mutations in genomes. However, the success rate of incorporation of the desired mutations at the target site is difficult to predict and therefore must be empirically determined. Here, we adapted the widely used TIDE method for quantification of templated editing events, including point mutations. The resulting TIDER method is a rapid, cheap and accessible tool for testing and optimization of template-directed genome editing strategies.

---

The CRISPR system for genome editing has become one of the most popular techniques in molecular biology. CRISPR endonucleases such as Cas9 can cleave genomic DNA with high precision, and due to error-prone repair mechanisms this can result in small insertions or deletions (indels)^1-3^. Alternatively, precisely designed small nucleotide changes can be incorporated near the break site by providing a donor template^4, 5^, such as a single-stranded oligodeoxynucleotide (ssODN)^4, 6^. By homology-directed repair (HDR), the DNA of the donor template is exchanged with the genomic DNA, and thereby the desired mutations are introduced^7, 8^. Such precise editing offers the possibility to create and study specific mutations, or to correct disease-causing nucleotide variants^5, 9^.

A current limitation of this template-directed strategy is that the efficacy is unpredictable and often low. Because error-prone non-templated repair pathways are active besides HDR, various indels are often introduced at the target site instead of the desired mutation. Moreover, a substantial fraction of the target sequence may remain unaltered. Thus, exposing a pool of cells to CRISPR and a donor template yields a complex mixture of cells with wild-type DNA, indels and the designed mutation, with unpredictable ratios^10-12^. A quick and easy assay to determine these ratios is of key importance, particularly if one wants to estimate how many cells are to be cloned from the pool in order to obtain at least one clonal line with the desired mutation.

High throughput sequencing of DNA around the induced break site is a powerful tool to analyze the mutation spectrum^13^, but is also expensive and requires substantial computational analysis. The frequently used Tracking of Indels by DEcomposition (TIDE) method^14^ is much simpler and cheaper, as it requires only two standard Sanger capillary sequencing reactions and an easy-to-use web tool for data analysis. However, in its present form TIDE is not suitable for templated genome editing, because it can only detect overall indel frequencies and not nucleotide substitutions or specifically designed indels. Here, we present TIDER (Tracking of Insertions, DEletions and Recombination events), a modified version of TIDE that estimates the incorporation frequency of any type of template-directed mutations, together with the background spectrum of additional indels. The corresponding TIDER web tool is freely accessible at http://tide.nki.nl.

The original TIDE protocol requires two capillary sequencing traces from a DNA stretch around the editing site: one test sample (DNA from cells treated with targeted nuclease) and one control (e.g. DNA from mock transfected cells). In addition, a text string representing the sequence of the sgRNA is used as input to determine the expected break site. Indels are then quantified by computational decomposition of the mixture of sequences in the test sequence trace, using the control sequence trace for comparison^14^.

TIDER takes a similar decomposition approach, but it requires one additional capillary sequencing trace. This “reference” trace is derived from a pure DNA sample that carries the designed base pair changes as present in the donor template. Such a reference trace can be generated readily from commercially synthesized DNA or from DNA obtained by a simple two-step PCR procedure as outlined in **Supplementary Figure S1** and Methods. The latter approach requires slightly more hands-on time, but is typically quicker and cheaper. Sequence traces derived from either source performed equally well in TIDER (see below).

To determine the individual sequence variants in the DNA of a cell pool, the algorithm decomposes the sequence trace of the experimental sample by multivariate non-negative linear modeling (**Figure 1**). For this, it uses the control and reference traces to construct a set of models of all likely outcomes of the cutting and repair process: wild-type sequence, all possible random indels at the break site, and the desired sequence as result of HDR. All of these models are collectively fitted to the experimental sample trace. The software provides an R^2^ value as a goodness-of-fit measure, and calculates the statistical significance of the detected HDR events. Additionally, TIDER generates a set of quality control plots that enable the user to verify the expected break site, and to visually inspect the sequence changes resulting from the editing process (**Supplementary Figure S2**).

**Figure 1.**
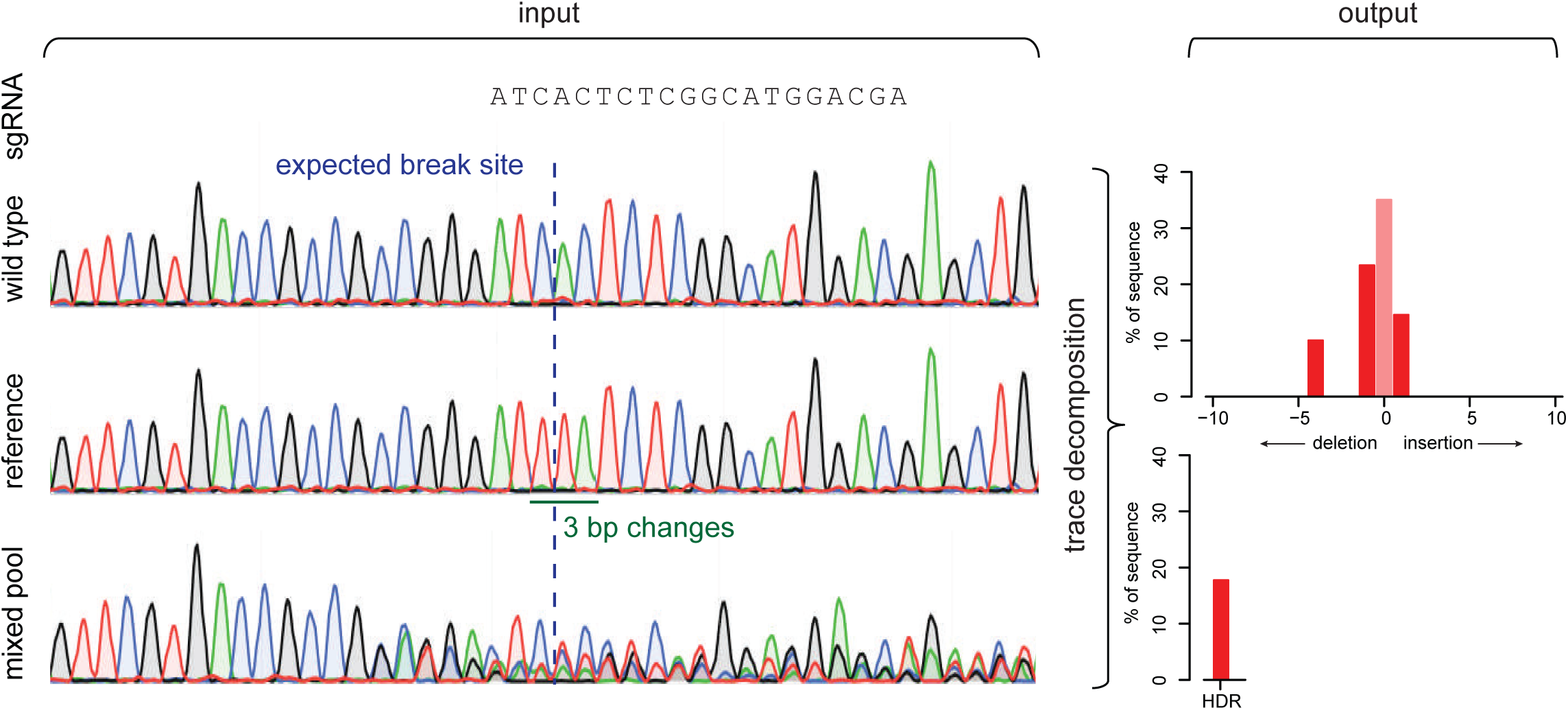
Assessment of homologous direct repair by sequence trace decomposition. Overview of TIDER algorithm and output. The introduction of designed mutations by homology directed repair with a donor template results in defined changes in a sequence trace. Due to NHEJ repair also insertions and deletions arise at the targeted break site. All these mutations yield in a composite sequence trace. As input a sgRNA sequence string and three sequences are required: 1) wild-type control, 2) reference file with designed mutations in the used donor template and 3) composite test sample. Trace decomposition yields the spectrum of indels and the HDR events with their frequencies (See main text and http://tide.nki.nl for explanation).

To test the performance of TIDER, we initially mimicked the occurrence of HDR events *in vitro* by mixing DNA carrying defined sequence variants. First, we combined “wild-type” DNA with “mutant” DNA carrying a single base pair change in various ratios. We performed standard capillary sequencing and analyzed the resulting data with the TIDER software. The algorithm was able to detect the single base pair change quantitatively with a sensitivity down to ~5% at a p-value cutoff of 0.01 (**Figure 2a, b** and **Supplementary Figure S3a**). Only very small amounts of false-positive indels were scored across the entire range of mixing ratios (**Supplementary Figure S4**). No statistically significant signal was detected when a reference sequence with a different point mutation was used, attesting to the specificity of TIDER for one particular mutation (**Figure 2b,** purple triangles). More complex mixtures consisting of wild-type DNA, DNA carrying various indels and DNA with a single base pair change could also be resolved accurately (**Figure 2a, c; Supplementary Figure S5**). In this particular experiment the proportion of the designed mutant was somewhat overestimated at low mixing ratios, but with increasing ratios the estimates were accurate. Results were nearly identical for reference DNA generated by full synthesis or by the two-step PCR procedure (compare **Figure 2b-c** and **Supplementary Figure S3b-c**). In another mixing experiment with a different complex pool and a different mutant, the accuracy was substantially higher (**Supplementary Figure S3d-f**), presumably because this mutant differed at four base pair positions from the wild-type DNA instead of one position.

**Figure 2.**
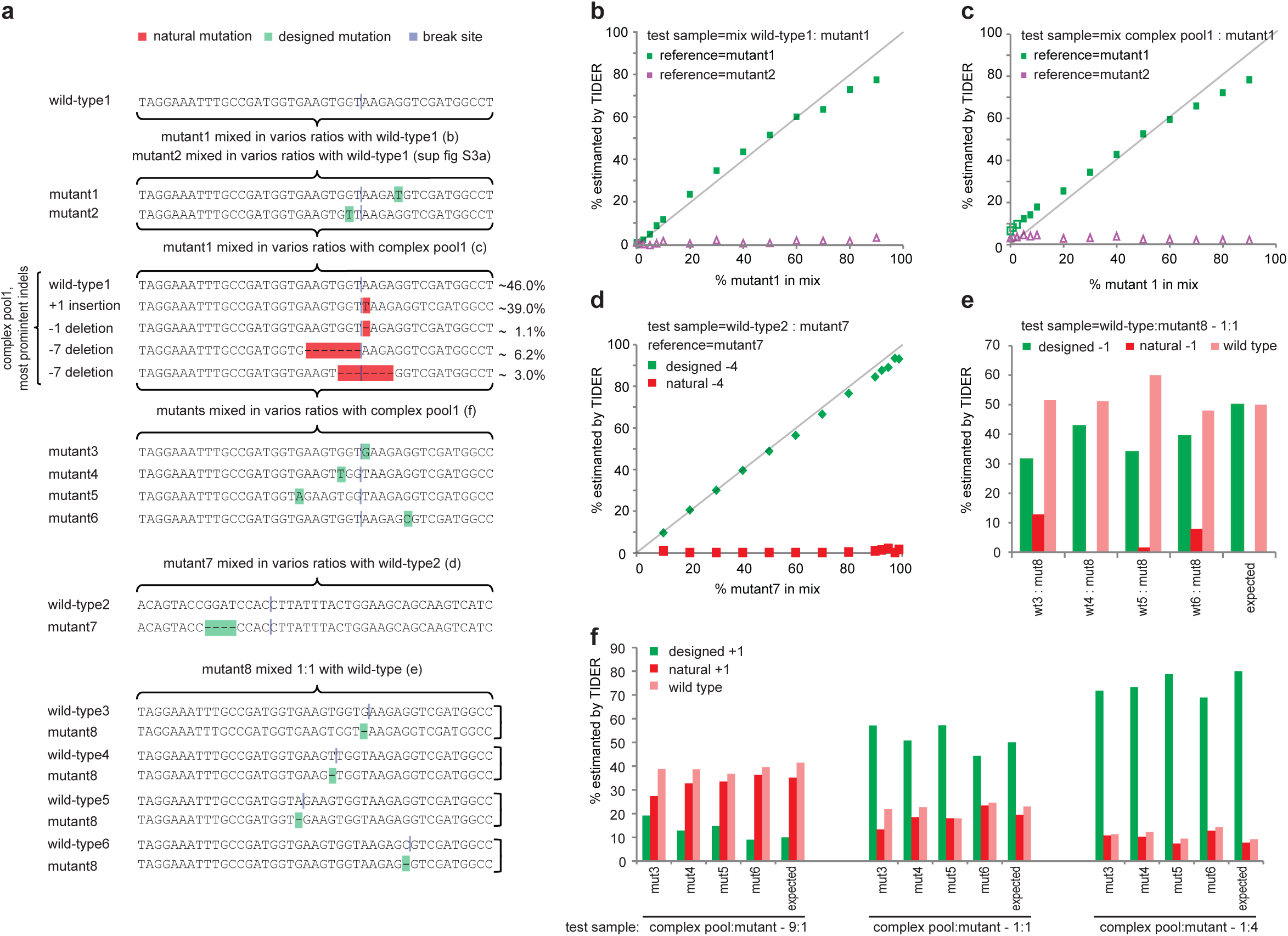
TIDER decomposition of *in vitro* mixes of DNA. Template-directed genome editing experiments in a pool of cells were simulated by *in vitro* mixing of DNA fragments carrying specific mutations with a corresponding wild-type DNA fragment,or with a complex pool of DNA fragments carrying different indels. **(a)** DNA mixtures that were tested. Letters in parentheses refer to the panels that show the corresponding TIDER results. Only the relevant sequences of the tested DNA fragments are shown; the total length of the fragments was 529 bp. “Designed” mutations are indicated in green, “natural” indels in red. Virtual Cas9 break sites used in these analyses are marked in dark blue. The complex pool is DNA from a pool of cells treated with Cas9 and sgRNA; it contains wild-type DNA as well as various indels introduced by NHEJ, of which the relative amounts are indicated. **(b-c)** PCR product with mutation1 was mixed in indicated relative amounts (horizontal axis) with wild-type DNA or with the complex pool. The proportion of mutant DNA was determined by TIDER (vertical axis) using either correct reference (mutant1, green squares) or incorrect reference (mutant2, purple triangles). See **Supplementary Figures S4 and S5** for the complete decomposition results. **(d)** Same as (b-c), but for wild-type2 mixed at various ratios with mutant7 that carries a -4 deletion.Green diamonds: estimated “designed” -4 deletions as in the reference file. Red squares: estimated “natural” -4 deletions (i.e. all deletions of size 4 that overlap with or are immediately adjacent to the break site). **(e)** 1:1 mixtures of mutant8 and wild-type3-6. For the TIDER analysis mutant8 was used as reference and the respective break sites were chosen as indicated in (a); hence in each analysis mutant8 carries a “designed” -1 deletion relative to the wild-type DNA. The percentages of the designed -1, natural -1 (other deletions of size 1) and wild-type DNA as estimated by TIDER are shown. The expected percentages are depicted in the last column. **(f)** TIDER analyses of mixtures of the complex DNA pool with each of mutant3-6 at three different ratios (9:1, 1:1, and 1:4). Bar graphs show percentages of the designed +1, natural +1 (other insertions of size 1) and wild-type DNA as estimated by TIDER. Expected percentages are depicted in the last column of each mixture set. In all analyses in (**b-f**) default TIDER settings were used (size range 0-10 for deletions and 0-5 for insertions).

A potentially more challenging scenario is when the templated mutation is a small deletion. During the repair process, other (non-templated) deletions of the same size may arise. We tested the ability of TIDER to discriminate the designed deletion from alternative deletions of the same size. When we mixed wildtype DNA with varying amounts of a -4 deletion, TIDER correctly determined the deletion with high specificity as “designed” when DNA carrying this deletion was used for the reference trace (**Figure 2a, d**). Similar results were obtained with four different “designed” -1 deletions, although in two instances a small fraction was scored as non-templated deletion (**Figure 2a, e**). Therefore, in the presence of only a small designed deletion (-1, -2) near the expected break site the designed mutation may be underestimated somewhat. In general, however, TIDER does not mistake a “designed” deletion for a non-templated deletion of the same size.

As a more stringent *in vitro* test, we generated several mutant sequences with a +1 insertion at various positions relative to the break site (**Figure 2a**), and mixed each of these “designed” mutant DNAs with a complex pool of DNA that contained ~39% of “natural” +1 insertions. TIDER analysis resolved the composition of the mixtures with high accuracy (**Figure 2a, f**). Sequencing of the opposite DNA strand yielded very similar results (**Supplementary Figure S6**), illustrating the robustness of the approach. This experiment illustrates that the presence of a non-templated insertion generally does not compromise the detection of the designed insertion of the same size. Together, these *in vitro* mixing experiments show that sequence trace decomposition can in most cases accurately identify and quantify “designed” mutations (base pair substitutions as well as small deletions and insertions) in a complex background of indels caused by imperfect repair.

Next, we tested TIDER in a series of *in vivo* experiments in which we subjected specific genomic sequences to templated editing in mouse embryonic stem (mES) cells and human retinal pigment epithelium (RPE-1) cells. We co-transfected these cells with Cas9, a sgRNA and a corresponding ssODN carrying 3 or 4 nucleotide substitutions. As control the ssODN was omitted. To verify the TIDER results, we sequenced the same samples by next generation sequencing (NGS). For 5 out of 5 tested sgRNA/ssODN combinations we found that the NGS results are similar to the TIDER estimations (**Figure 3a-e**). Moreover, in cells treated with sgRNA in the absence of a donor template, TIDER detects almost no HDR events, while the non-templated indel spectra are again highly similar to those determined by NGS (**Figure 3f & Supplementary Figure S7**). Furthermore, in one set of editing experiments involving a complex set of templated nucleotide substitutions, application of TIDER with different window settings, combined with the data visualization tool, reproducibly revealed that one nucleotide substitution 40bp upstream of the break site was less efficiently incorporated than the more proximal substitutions; this result was confirmed by NGS (**Supplementary Figure S8**). We conclude that TIDER can reliably estimate the frequency of HDR events in a background of non-templated indels in genomic DNA from pools of cells.

**Figure 3.**
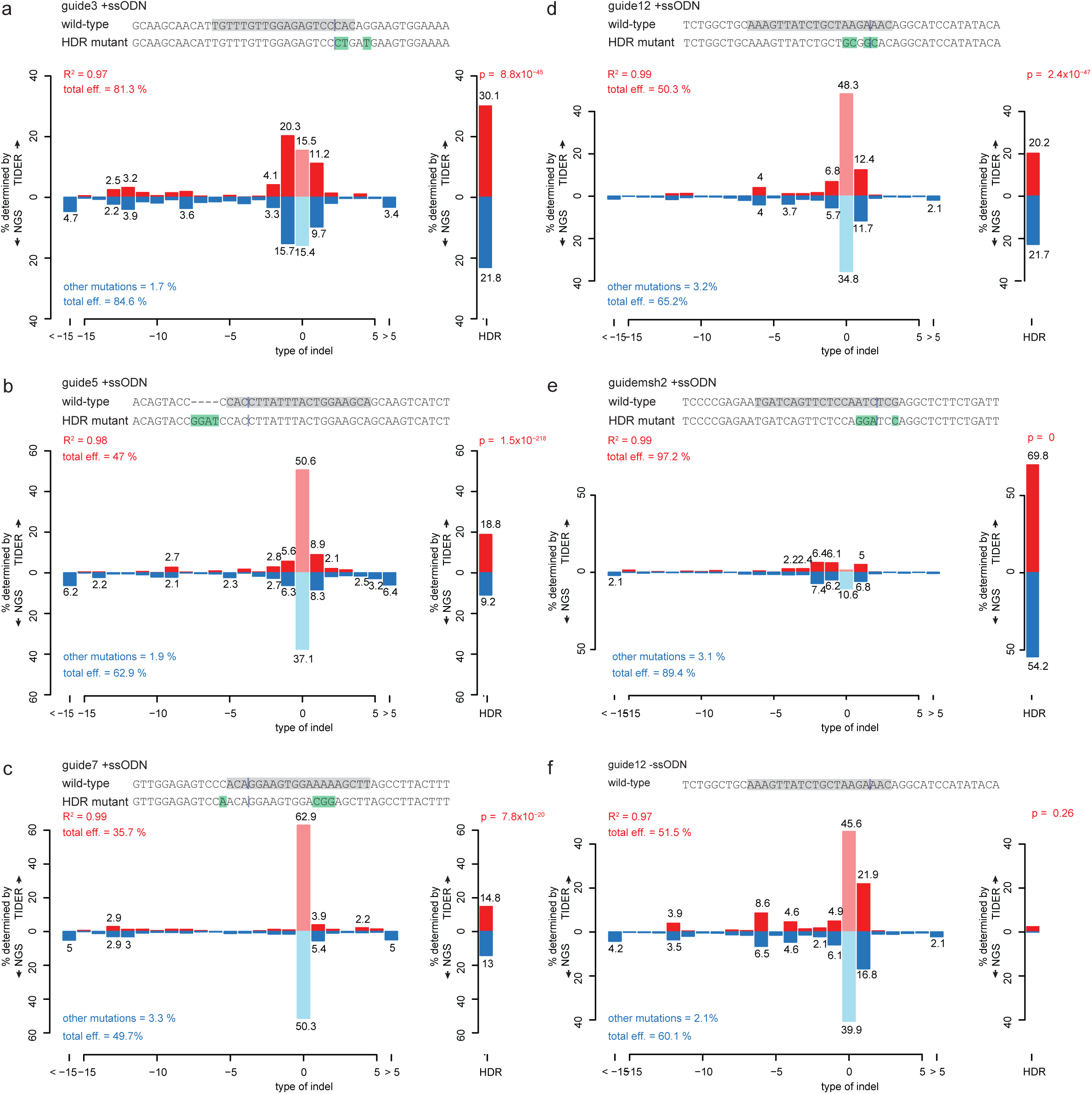
Application of TIDER to *in vivo* edited DNA sequences. Comparison of TIDER and NGS analyses of various mutations introduced by template-directed Cas9 editing in human cell line RPE **(a-d,f)** and mouse ES cells **(e)**. In each panel (a-e), a pool of cells was treated with Cas9, a targeting sgRNA and a ssODN carrying 3-4 mutations. Panel (**f**) shows a control experiment corresponding to (**d**) in which the ssODN was omitted. Additional control experiments corresponding to (**a-c**) are shown in **Supplementary Figure S7**. In each panel, the top sequence corresponds to wild-type, with the sgRNA sequence highlighted in grey and the expected cut site marked by a vertical line; the bottom sequence indicates the designed mutant, with mutated nucleotides highlighted in green. Bar graphs show the estimated percentage of successfully edited DNA molecules (right-hand plot; “HDR”) and of indels of the indicated size (left-hand plot). Upward axes show TIDER estimates; downward axes show the NGS estimates based on the same DNA sample. Pale red and blue bars indicate proportions of wild-type (non-mutated) sequence. R^2^ values indicate the goodness-of-fit score for the TIDER estimates; “total eff” indicates the total according to TIDER (top) and NGS (bottom); “other mutations” are all non-indel, non-designed mutations as detected by NGS (and which cannot be detected by TIDER). For TIDER, the decomposition was limited to deletions of sizes 0-15 and insertions of sizes 0-5. For NGS, at least 2×10^4^ reads were analyzed in each experiment.

In summary, TIDER is a simple and rapid assay to evaluate the efficacy of templated editing. Like TIDE, TIDER requires only standard capillary sequencing, thereby offering a widely accessible, cheap and rapid alternative to NGS. TIDER is much more quantitative and informative than the Surveyor and T7 endonuclease I cleavage assays^15^ ^16^, which are unable to discriminate between the designed mutation and randomly induced indels.

TIDER is primarily designed to determine the efficacy of templated genome editing. It complements TIDE, which can only detect non-templated indels. While TIDER can also quantify the latter, TIDE is more suitable for the assessment of non-templated editing experiments because it is slightly simpler in experimental design. Both web tools are freely available through http://tide.nki.nl/.

Because the TIDER algorithm analyses individual peak heights in the input sequence traces, the accuracy of TIDER relies on the quality of the PCR products and the sequence reads. This is particularly relevant when the difference between the wild-type and reference sequence is small, e.g., in case of single-nucleotide differences. In such cases we recommend that the results are verified by sequencing of the opposite strand. The TIDER web tool provides graphical feedback as well as an R^2^ value as means to estimate the reliability of the analysis. We generally recommend that R^2^ is above 0.9. While the default settings of the web tool are suited for most purposes, parameter settings can be adjusted interactively to optimize the performance. We also recommend that results are verified by sequencing of the opposite strand. Note that the TIDER algorithm cannot resolve HDR events that have acquired an additional non-templated indel, but the frequency of such double templated/non-templated mutations has been reported to be low when a PAM disrupting mutation is introduced^17, 18^.

## Methods

### Cell culture and transfection

Human retinal pigment epithelial (hTERT RPE-1, ATCC CRL-4000) cells were cultured in DMEM (Gibco 31966) supplemented with 10% fetal bovine serum (FBS, HyClone^®^). Mouse embryonic stem cells (mESCs) were cultured as described^19^. Briefly, mESCs were expanded and maintained on sublethally irradiated mouse embryonic fibroblast feeder cells in LIF supplemented medium. Prior to transfection, cells were seeded on gelatin-coated plates and cultured in Buffalo Rat Liver cell (BRL) conditioned medium supplemented with LIF (ESG1107, Merck (Millipore)).

The desired mutations were introduced in hTERT RPE-1 according to the RNP CRISPR approach of IDT. The sgRNAs were designed using CRISPR design tools of Benchling or MIT tool^20^. In brief, 1×10^5^ cells were seeded out the day before transfection in 12-well dish in 750 µL medium with 1 µM final concentration DNA-PKcs inhibitor NU7441 (Cayman). 3 µL of 10 µM sgRNA and 3 µL of 10 µM Cas9 protein were mixed in optiMEM (Life Technologies) to final volume of 125 µL and incubated in for 5 min at RT. 4.5 µL of this sgRNA/Cas9 mix, 1.5 µL of 10 µM ssODN (Ultramer IDT) and 4.5 µL Lipofectamine RNAiMAX (Invitrogen) were added to 240 µL optiMEM. Mixture was incubated at RT for 20 minutes before adding to the cells. The next day the medium was changed, and 2 days after transfection the cells were harvested for analysis of the genomic DNA.

mESCs were seeded 2 days before transfection at a density of 5×10^4^ cells in each well of a 6-well dish. 250ng of a PX330 derived vector (Addgene #42230, with an added puromycin resistance cassette) and 2.25µg of ssODN were added to 250µL optiMEM. 6.25µl of Mirus TransIT LT-1 was added to this mixture and mixed by pipetting. After incubation for 15 minutes at RT, the solution was added dropwise to the cells. One day after transfection, the cells were reseeded on gelatin coated plates in BRL medium containing 3.6µg/mL puromycin. 2 days after reseeding, the medium was replaced without puromycin, and 4 days later the cells were harvested for genomic DNA extraction.

The following sgRNA sequences were used:

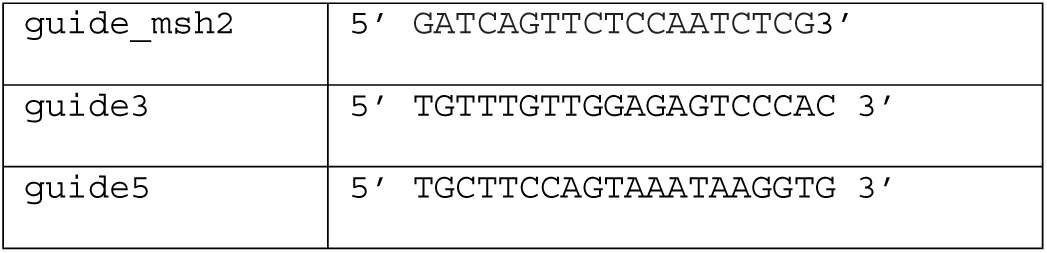

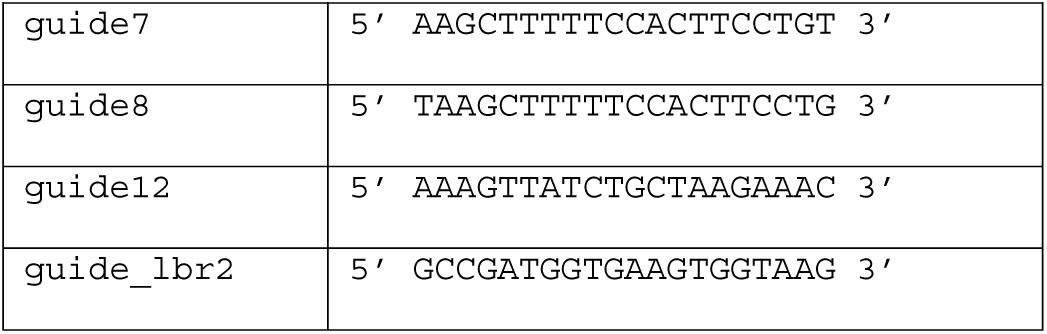

The following ssODNs sequences were used:

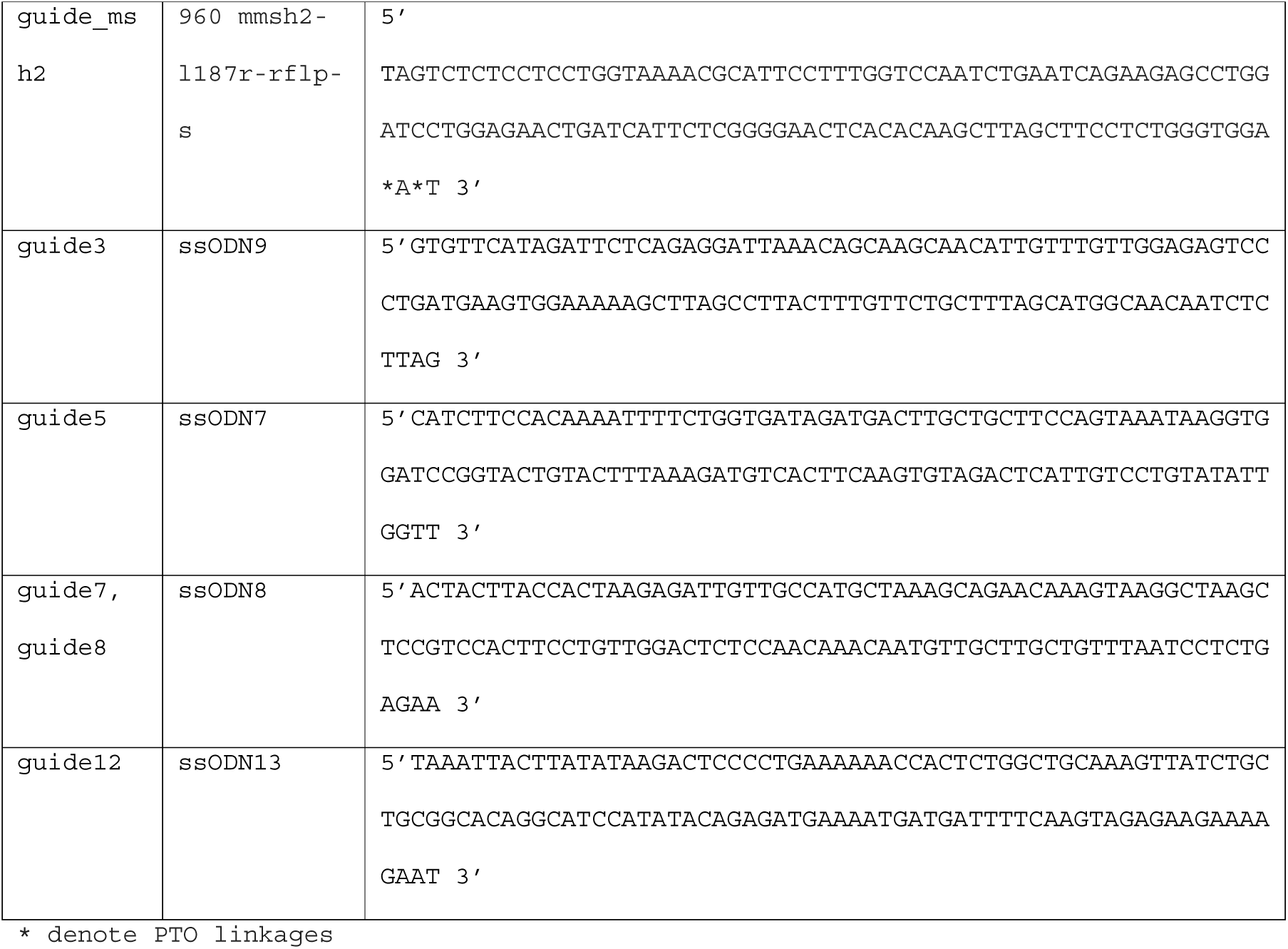

### PCR control & test sample

Genomic DNA was isolated 2 days (RPE cells) or 7 days (mESC cells) after transfection using either the Isolate II Genomic DNA Kit (Bioline) or lysisbuffer (100mM Tris (pH 8.5), 50mM EDTA, 40mM NaCl, 0.2%SDS and 100ug/mL proteinase K) for 2 hours at 55°C followed by 45 minutes incubation at 85 °C and DNA precipitation by addition of 2.5 volumes of 100% ethanol, followed by 30 minutes centrifugation at 14,000 RPM at 4 °C. After washing of pellets with 70% ethanol, pellets were dissolved in TE buffer by overnight incubation at 55 °C. PCR reactions were carried out with 50 ng genomic DNA in MyTaq^TM^ Red mix (Bioline) according to manufacturer’s instructions using primers a & b (10 µM) as listed below. PCR thermocycling scheme: 1 min at 95°C (1×), followed by 15 sec at 95°C, 10-20 sec at 55-60°C, and 10-20 sec 72 °C (25-35×). The PCR products were purified using the PCR ISOLATE II PCR and Gel Kit (Bioline). The Msh2 target site was amplified with Taq polymerase (MRC Holland) using the following PCR program: 2 min 94 °C (1×), followed by 30 sec at 94 °C, 3 sec at 53.8 °C and 40 sec at 72 °C (37×) and 5 min at 72 °C (1×).

The following primer pairs spanning the target site were used (a: forward; b: reverse):

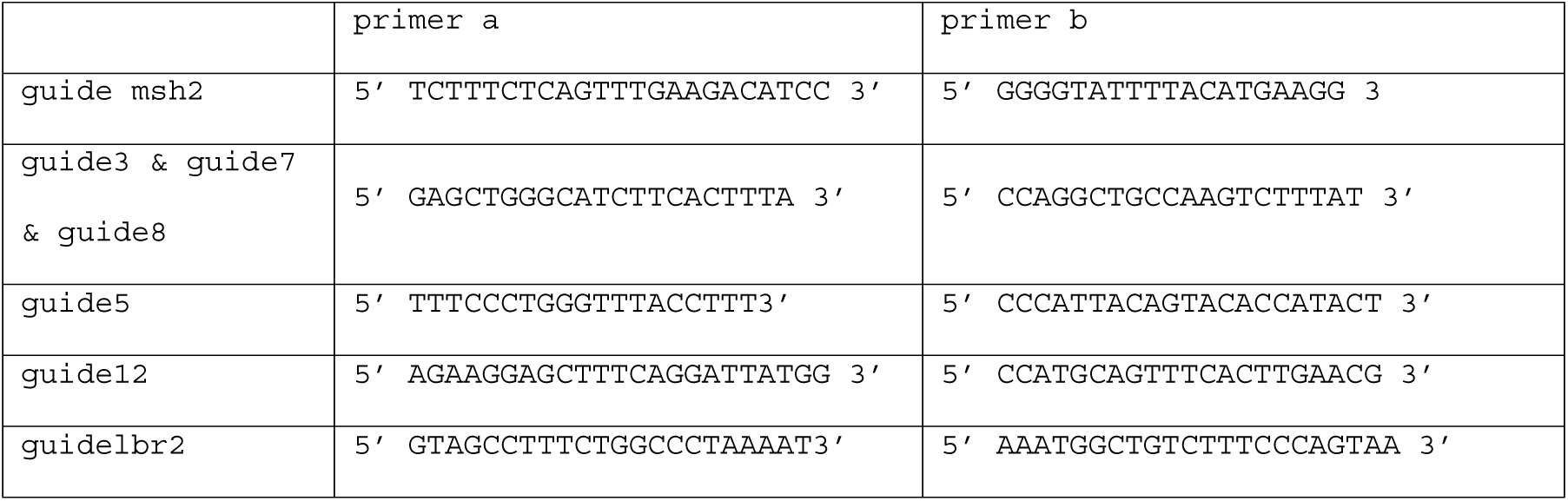

### PCR Reference sample

The reference sequence was generally generated in a 2-step PCR reaction (**Supplementary Figure S1**). Two complementary primers (primers c & d) were designed that carried the designed mutations as present in the donor template. Two standard PCR reactions were done with 50 ng wild-type genomic DNA in MyTaq^TM^ Red mix (Bioline) using primers a & c and primers b & d. PCR thermocycling scheme: 1 min at 95°C (1×), followed by 15 sec at 95°C, 15 sec at 55°C, and 20 sec 72°C (25-30x). The two PCR products were purified using the PCR ISOLATE II PCR and Gel Kit (Bioline). Next, the resulting two PCR amplicons (each 1 µL) were combined with 48 µL buffer (10 mM Tris, 50 mM NaCl, 1 mM EDTA) and denatured for 5 min 95 °C and cooled down (0.1 °C/sec) to 25 °C. Of this mixture 3 µL was subsequently used as template in a PCR reaction with MyTaq^TM^ Red mix (Bioline) with primers a & b, starting with an extension step as follows: 15 sec at 72°C (1×), followed by 15 sec at 95°C, 15 sec at 55°C, and 20 sec 72°C (25-30x). The PCR products were purified using the PCR ISOLATE II PCR and Gel Kit (Bioline).

The following primer pairs spanning the to be edited site were used (c: reverse; d: forward):

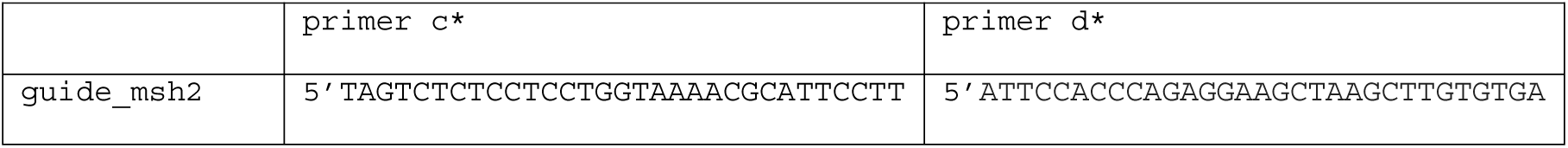

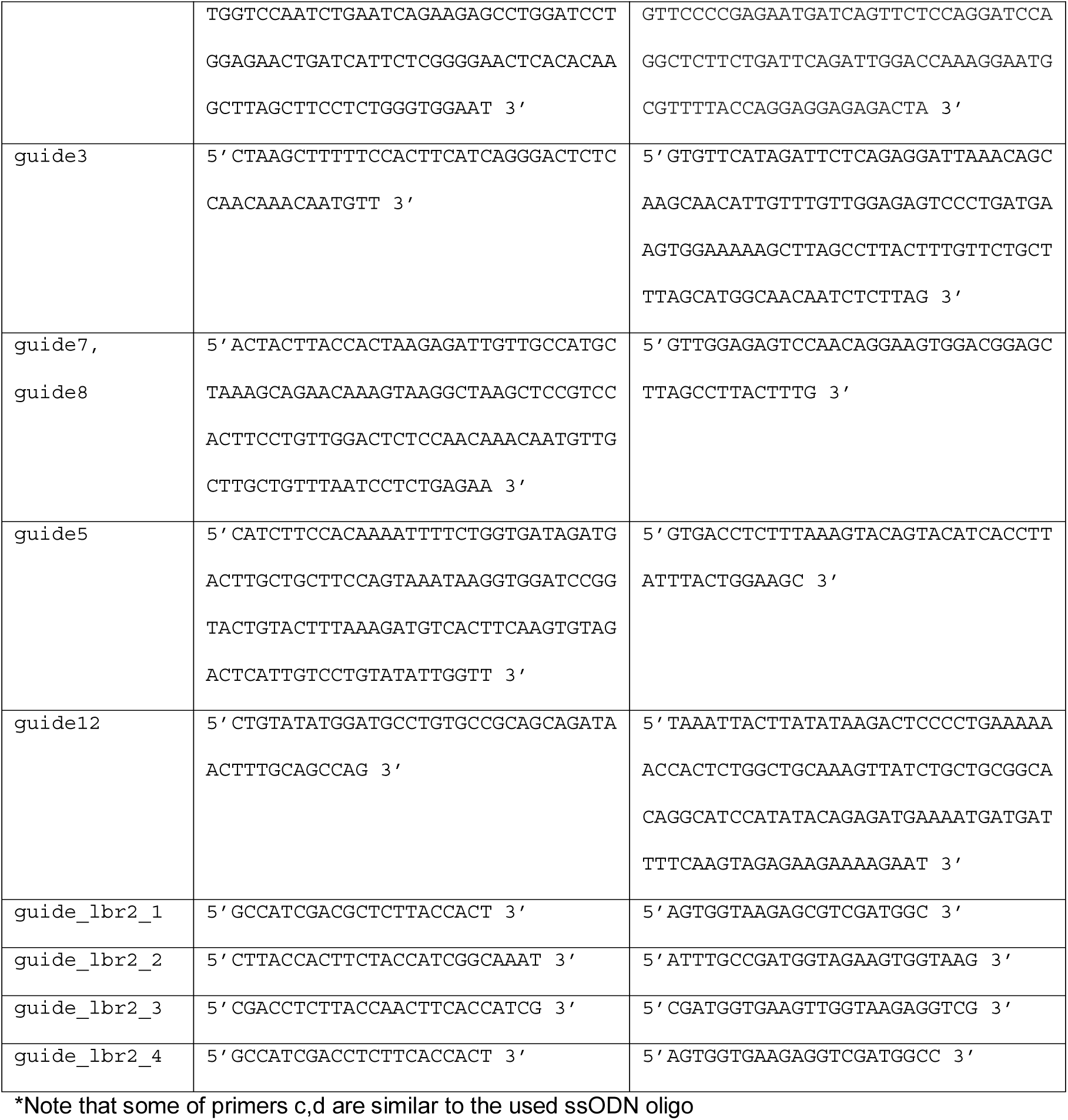

### Sanger sequencing

Purified PCR samples (100 ng) were prepared for sequencing using 4 µL of BigDye^®^ terminator v3.1 (Applied Biosystems^®^) and 5 pmol primer in final volume of 20 μl. Thermocycling program: 1 min at 96°C (1×), followed by 30 sec at 96°C, 15 sec at 50°C, and 4 min at 60°C (30×), and finishing with 1 min incubation at 4°C (1×). Sequence traces were generated on an Applied Biosystems 3730xl DNA Analyzer, running 3730 Series Data Collection Software V4 and Sequencing Analysis Software V6.

### Next generation sequencing

PCR was performed in two steps with genomic DNA as template; PCR1 with ~50 ng genomic DNA and site specific barcoded primers. PCR2 used 2 µL of each PCR1 product with Illumina PCR Index Primers Sequences 1–12. Each sample was generated with a unique combination of a barcode and index. Both PCR reactions were carried out with 25 µL MyTaq Red mix (Bioline), 4 µM of each primer and 50 µL final volume in a 96 well plate. PCR conditions were 1 min at 95 °C, followed by 15 sec at 95 °C, 15 sec at 58 °C and 1 min at 72 °C (15x). 20 µL of 8 samples were pooled and 100 µL was loaded onto a 1% agarose gel. PCR product was cut from gel to remove the primer dimers and cleaned with PCR Isolate II PCR and Gel Kit (Bioline). The isolated samples were sequenced by Illumina MiSeq.

The following primers sequences were used for NGS:

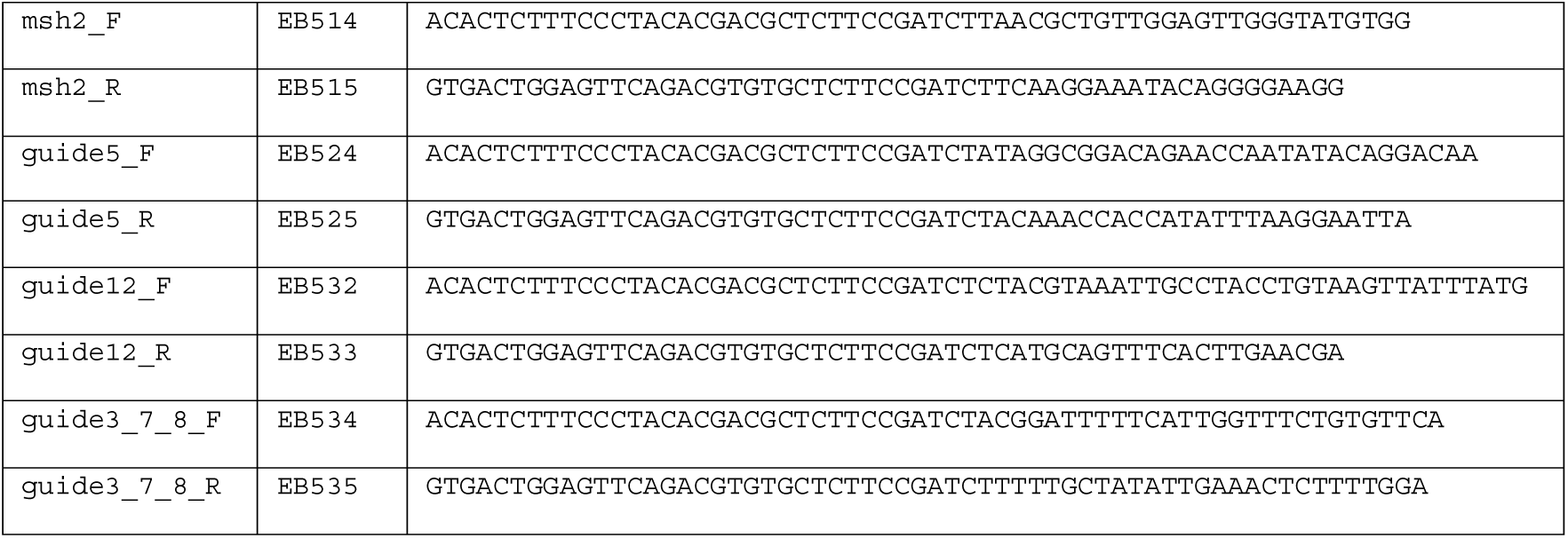

### NGS data analysis

In order to identify insertions and deletions, the distance between a fixed sequence ~50 nt upstream of the break site and ~50 nt downstream of the break site was determined. *Insertions* and *deletions* have a distance longer or shorter than wild-type, respectively. For each of the remaining reads a window of 50 nucleotides (from -25 to +25 relative to the expected break site) was compared to the corresponding nucleotide sequence strings of the control and reference sequences. Windows with zero or one mismatches compared to the control sequence were counted as *wild-type* reads. Subsequently, remaining reads with zero or one mismatches compared to the reference sequence were counted as *HDR reads*. All other reads are counted as *other mutations*. Reads in which we could not find a match with the constant parts are discarded. Finally, for each sample, the ratio of each mutation type over the total of reads is calculated.

### TIDER software

TIDER is built upon the previous published TIDE software^14^. TIDER code was written in R, version 3.3.2. TIDER requires as input a control sequence trace file (e.g. obtained from cells transfected without Cas9), a sample sequence trace file (e.g. DNA from a pool of cell treated with Cas9 and donor template), a reference sequence trace file (e.g. DNA from the donor template) and a character string representing the sgRNA sequence (20 nt).

We advise to sequence a stretch of DNA ~700 bp enclosing the designed editing site. The projected break site should be located preferably ~200 bp downstream from the sequencing start site. The sequencing data files (.abif or .scf format) are parsed using R Bioconductor package *sangerseqR*^21^ (version 1.10.0). Additional parameters have default settings but can be adjusted if necessary. The web interface was constructed using the *shiny* R package (version 1.0.0).

Briefly, the algorithm consists of the following steps. Both the test sample and the reference sequence are first aligned to the control sample sequence using standard Smith-Waterman local alignment implemented in the *BioStrings* package (version 2.42.1) in Bioconductor^22^. Subsequent calculations are done using the peak heights of the four bases for each position in the aligned sequence trace data. Next, for each position, the absolute peak height of each base is converted to a relative peak height by dividing it by the sum of the peak heights of all four bases at that position. All subsequent calculations are done using these relative peak heights.

In contrast to TIDE, the decomposition window of TIDER spans by default from 20bp upstream of the break to 80 bp downstream from the break. This window can be interactively adjusted, but it should contain all nucleotides that are edited. Within this window, sequence trace models are constructed of all possible indel occurrences that may realistically be expected, i.e, deletions of sizes {0…n} and insertions of sizes {0…m} that overlap with or are immediately adjacent to the break site. For example, to model all possible -4 deletions, 5 different sequence trace models are constructed; and to simulate all possible insertions of size 3, 4^3^ = 64 trace models are constructed. By default, n is set to 10, and m to 5. For deletions, the model traces are simply constructed from the control trace by deleting the values at the corresponding positions. For insertions, the average value of the same nucleotide occurrence within the whole sequence trace is used. The break site is assumed to be between the 17th and 18th bases in the sgRNA target sequence (3 bp before the PAM)^23^. The sequence trace models are constructed accordingly for each of the four bases, after which the vectors of the four bases are concatenated, so that each model consists of a single vector. Subsequently, control sequence model, all indel models and the reference sequence model are combined into a single decomposition matrix. To avoid doublet models, in case the reference consists of an insertion or deletion at the break site, the identical simulated insertion or deletion is removed from the decomposition matrix.

The decomposition is subsequently performed in two iterations. First, the sequence trace from the test sample is assumed to be a linear combination of the wild-type trace, the modeled indel traces and the reference trace. This combination is decomposed by standard non-negative linear modeling, for which we used the R package *nnls* (version 1.4). After this first trace decomposition, all sequence variants with an estimated frequency of exactly 0 are removed, and the decomposition is repeated with the remaining models.

Next, the frequencies of the various traces of same deletion or insertion size are summed. R^2^ is calculated to assess the goodness of fit. The p-value associated with the estimated abundance of the reference trace is calculated by a two-tailed t-test of the variance-covariance matrix of the standard errors. Finally, the fitting coefficients (frequencies) are multiplied by a constant factor such that their sum equals R^2^.

### Plots for visual inspection of sequence traces

TIDER uses the relative peak heights to determine the abundance of aberrant nucleotides by subtracting the peak heights of the highest control nucleotide over the length of the whole sequence trace of either the test sample or reference. Then, the highest peaks in the reference and the peaks in the control at the same location that are not the highest are identified (the designed base pair changes). Of these positions the corresponding nucleotide peak signal in the control and test sample are plotted to show the relative incorporation of the donor template. The plots of these sequence signals allows the user to check the quality of the sequence data, inspect proper alignment, verify the expected cut site, and interactively select the region used for decomposition.

### TIDER settings and constrains

For TIDER, we have empirically determined an optimal decomposition window of 100 bp for most applications, but this can be interactively adjusted.

In case the designed mutation consists of an insertion larger than +1, TIDER does not consider natural insertions of the same size, because we found the decomposition to become less robust, and because we and others have rarely observed natural insertions larger than +1^14, 24^.

It has been reported that the incorporation of donor template is less efficient when the designed point mutations are further away from the break site^25^. This may confound TIDER estimates when such distal mutations are combined with mutations close to the break site. This is also what we observed (**Supplementary Figure S8**). By comparing different settings for the decomposition window and by visual inspection of the TIDER plots it is possible to infer such biases.

## Acknowledgement

We thank William Peters and Rubayte Rahman for assistance with setting up the web tool; NKI colleagues for software testing; members of the BvS lab for helpful suggestions.

## Author Contributions

EKB designed the study, performed experiments, wrote code, analyzed data, wrote the manuscript. ANK and TH performed *in vivo* editing experiments; CL and TC wrote code; JJ: supervised ANK; BvS supervised the study, wrote the manuscript.

